## Supplementary Information for "Extracellular phase separation mediates storage and release of thyroglobulin in the thyroid follicular lumen"

### Supplementary Figures

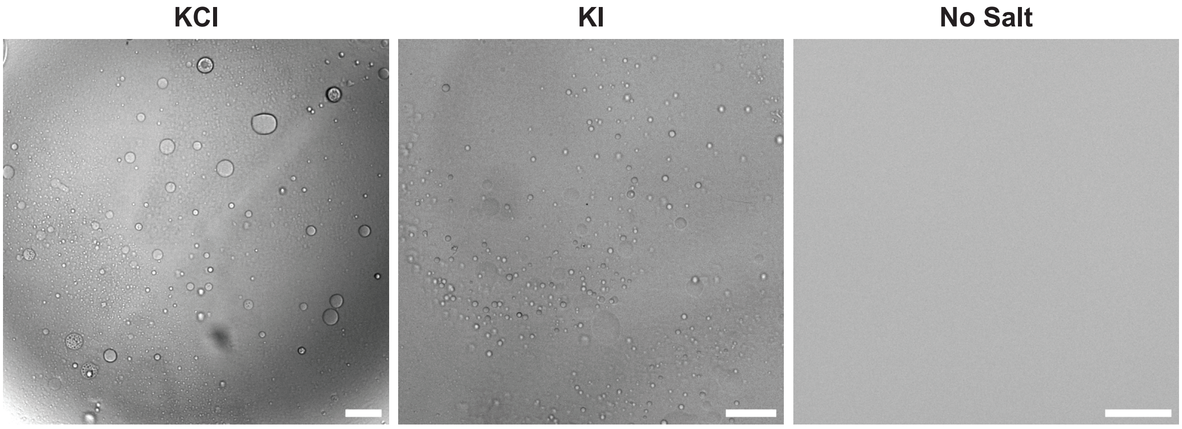


**Supplementary Figure S1. Unlabelled Tg forms condensates *in vitro*.** Left, unlabelled Tg condensates formed under 250 mM KCl. Middle, unlabelled Tg condensates formed under 250 mM KI. Right, the system remains mixed without adding salt. No Tg condensate formed without salt. All images acquired under bright field. Scale bars, 50 µm. All samples contained 10 µM unlabelled Tg, 4 w/v% PEG 20k in 50 mM HEPES buffer pH=7.5.

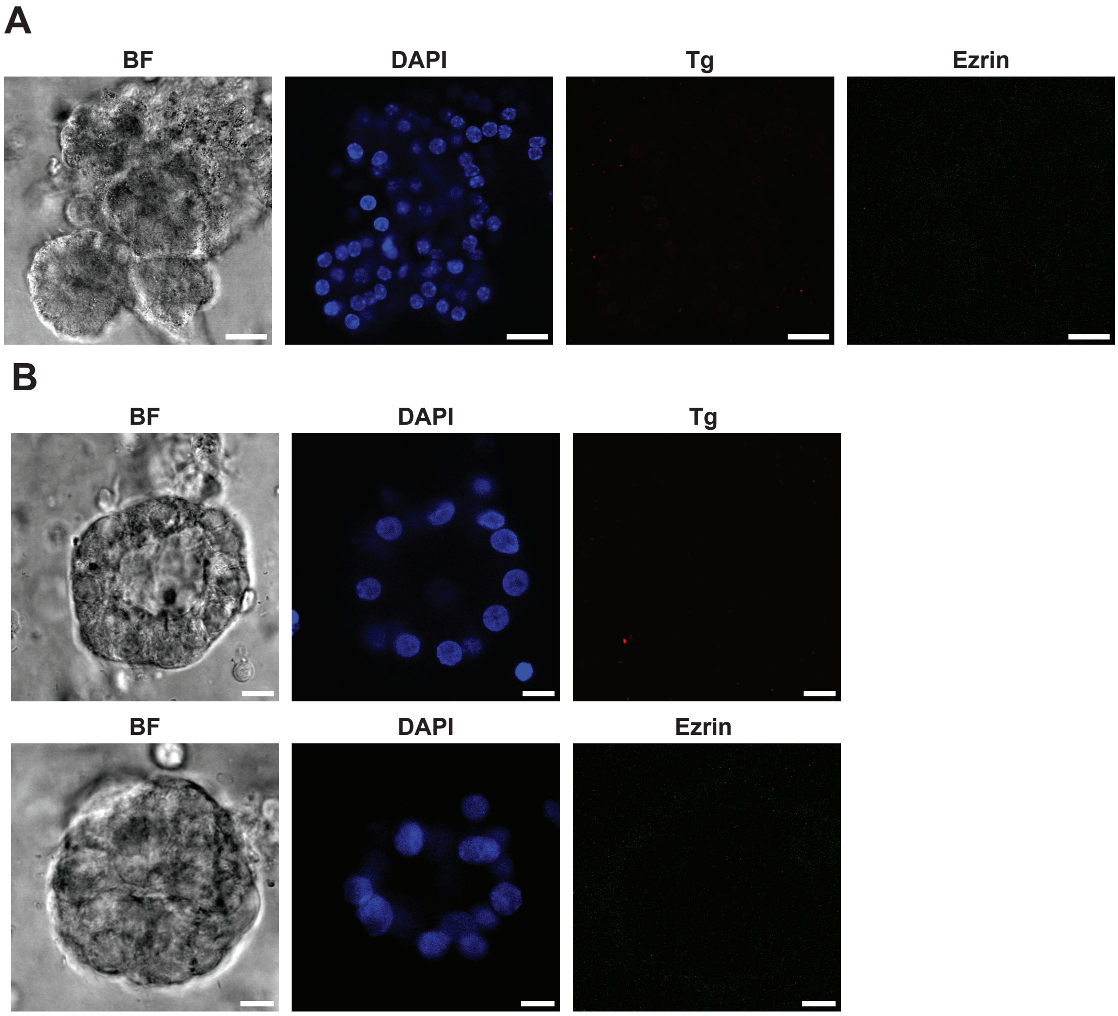


**Supplementary Figure S2. Isotype control images of Tg and ezrin in cultured mouse and human thyroid follicles.** Confocal images of Tg immunofluorescence isotype control experiment on **(A)** mouse follicles and **(B)** human follicles, imaged under bright field (BF), DAPI (thyrocyte nuclei), AF594 (Tg) and AF488 (ezrin) fluorescence channels. Follicles were observed under bright field; thyrocyte nuclei staining was also revealed under DAPI channel. No Tg or ezrin staining was observed, indicating high specificity of Tg and ezrin primary antibody binding. Scale bars, (A) 20 µm; (B) 10 µm.

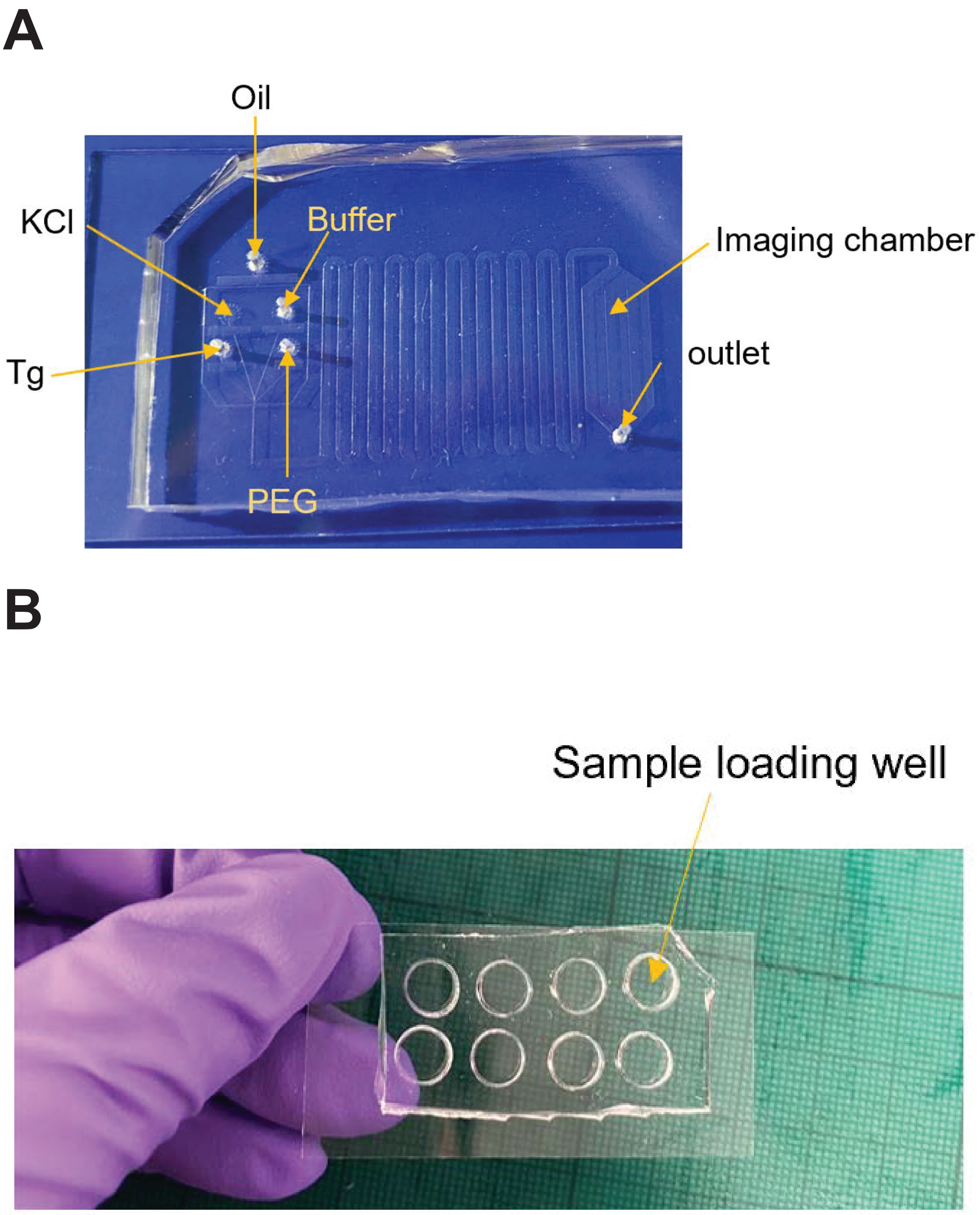


**Supplementary Figure S3. PDMS devices for PhaseScan experiments and imaging Tg condensates *in vitro*.** **(A)** The PDMS-based microfluidic device used for Tg PhaseScan experiment. This device has a droplet-making part containing 4 inlets for aqueous solutions and 1 inlet for oil; serpentine channels to stabilise oil flow rate and incubate droplets to a thermodynamically stable state before imaging, and an imaging chamber. **(B)** Sample loading well made of PDMS for confocal imaging. The chamber is made by PDMS covalently attached on glass coverslip.
